# α-syn overexpression, NRF2 suppression, and enhanced ferroptosis create a vicious cycle of neuronal loss in Parkinson’s disease

**DOI:** 10.1101/2022.07.22.501183

**Authors:** A Anandhan, W Chen, N Nguyen, L Madhavan, M Dodson, DD Zhang

## Abstract

Parkinson’s disease (PD) is the second most common neurodegenerative disorder, affecting millions each year. Most PD cases (∼90%) are sporadic, resulting from the age-dependent accumulation of pathogenic effects. One key pathological hallmark of PD progression is the accumulation of alpha-synuclein (α-syn), which has been shown to negatively affect neuronal function and viability. Here, using 3- and 6-month-old *Nrf2*^*+/+*^ and *Nrf2*^*-/-*^ mice overexpressing human α-syn (PD model), we show that loss of NRF2 increases markers of ferroptosis across PD-relevant brain regions. Increased ferroptosis was associated with an age- and genotype-dependent increase in α-syn pathology and behavioral deficits. Finally, we demonstrate that α-syn overexpression sensitizes neuronal cells and *ex vivo* brain slices to ferroptosis induction, which may be due to α-syn suppression of NRF2 at the protein level. Altogether, these results indicate that NRF2 is a critical anti-ferroptotic mediator of neuronal survival, and that the vicious cycle of α-syn overexpression and NRF2 suppression, leading to enhanced neuronal ferroptotic cell death, could represent a targetable and currently untapped means of preventing PD onset and progression.

## Introduction

The prevalence of Parkinson’s disease (PD) continues to increase as modern medical advances extend the average lifespan of the population. In the US alone, PD currently affects an estimated 1 million people each year, a number that is only expected to increase in the coming decades. Importantly, the estimated total cost of treating the ∼1 million US patients diagnosed with PD exceeded 51 billion dollars in 2017 and is projected to cost upwards of 79 billion to treat the ∼1.5 million patients diagnosed by 2037 [1]. While our understanding of the underlying causes of PD onset and progression has continued to increase over the years, treatment options remain limited, primarily focusing on alleviating symptoms. This is due in large part to the fact that by the time patients present in clinic with symptoms, much of the neuronal loss and glial dysfunction that drive PD pathogenesis has already occurred. Thus, a continuing goal in the field is the identification of pathological mechanisms that promote onset and early disease development, including therapeutic options that prevent neuronal loss, as opposed to just treating symptoms that appear once irreversible cell death has occurred.

One of the key pathological hallmarks of PD progression is the aberrant accumulation of α-synuclein (α-syn), which has been shown to have PD-relevant pathogenic effects in its monomeric, oligomeric, and insoluble fibrillar forms [2]. The notion that α-syn can be pathogenic across its different forms can have significant implications with regards to early versus late PD progression, as preventing aggregate formation or degrading aggregates that already exist, could significantly influence PD pathology. Critically, α-syn has been shown to enhance neuronal dysfunction and affect cell viability via a variety of mechanisms, including increased oxidative stress, mitochondrial dysfunction, altered membrane lipid dynamics, and autophagy dysfunction, among others [3]. A critical regulator of the response to these various stressors is the transcription factor nuclear factor erythroid 2-related factor 2 (NRF2) which regulates targets mediating redox, metabolic, and protein homeostasis [4]. Another critical function for NRF2 is preventing ferroptosis, a recently identified iron and lipid peroxide-driven form of cell death [5, 6]. Importantly, NRF2 has also been shown to decline with age [7], inferring that a gradual decline in its expression or function could be a significant contributor to PD onset and progression. This notion is supported by studies indicating that *Nrf2*^*-/-*^ mice exhibit greater PD pathology than their *Nrf2*^*+/+*^ counterparts, and that NRF2 activation is protective across a variety of PD contexts [8]. However, despite a clear understanding of the importance of NRF2 in preventing PD pathology, a strong understanding of what mode of cell death occurs remains unclear.

Along these lines, our previous study indicated that *Nrf2*^*-/-*^ mice overexpressing human α-syn (*SNCA*) exhibit worse α-syn-associated pathogenesis, dopaminergic neuron loss, and behavioral deficits at 3 months of age than their wild type counterparts [9]. However, despite our findings in this study, the exact cause of neuronal loss and behavioral deficits was not determined. Recently, several studies have shown that increased α-syn levels can induce ferroptosis in human iPSC-derived neurons with *SNCA* triplication [10, 11]. However, to our knowledge, there are currently no studies that have linked α-syn accumulation to ferroptosis induction *in vivo*. Intriguingly, while a key role for ferroptosis in neurodegeneration has been proposed across a variety of neurodegenerative disease contexts [12], very few studies have identified how and if ferroptosis occurs in translational models of PD. Despite this lack of experimental evidence, increased ROS, free iron, and lipid peroxidation, all of which are key hallmarks of ferroptosis, are also common features of brain regions affected during PD [13-15]. Accordingly, determining if ferroptosis occurs in relevant models of PD, as well as the role NRF2 plays, could represent a targetable and currently untapped mechanism of preventing PD onset and progression.

Here, using 3 and 6 month old *Nrf2*^*+/+*^ and *Nrf2*^*-/-*^ mice that overexpress human α-syn (*h*_α*-*_*Syn*^*+*^*:Nrf2*^*+/+*^ and *h*_α_*-Syn*^*+*^*:Nrf2*^*-/-*^), we show that loss of NRF2 increases markers of ferroptosis in the midbrain (MB) and striatum (ST) by 6 months of age. Increased ferroptosis was also associated with an age- and genotype-dependent increase in α-syn pathology and behavioral deficits. Finally, we demonstrate that _α_-syn overexpression sensitizes neuronal cells and *ex vivo* brain slices to ferroptosis induction, which may be due to _α_-syn-mediated suppression of NRF2 protein levels. Altogether, these results indicate that NRF2 is a critical anti-ferroptotic mediator of neuronal survival, and that α-syn may promote age-dependent neuronal loss and PD pathology via suppression of NRF2.

## Results

### Loss of NRF2 increases markers of ferroptosis by 6 months of age

To determine if genetic ablation of NRF2 has an effect on enhancing ferroptosis in the brain, the midbrain (MB) and striatum (ST) of 3 and 6 month old *h*_α_*-Syn*^*+*^*:Nrf2*^*+/+*^ and *h*_α_*-Syn*^*+*^*:Nrf2*^*-/-*^ mice were assessed for markers of ferroptosis. While there was no significant difference in the mRNA, and slight increase in the protein levels of *Ptgs2*/Cox2 or Acsl4 (pro-ferroptotic markers) between *h*_α_*-Syn*^*+*^*:Nrf2*^*+/+*^ and *h*_α_*-Syn*^*+*^*:Nrf2*^*-/-*^ mice at 3 months, both were elevated in the 6 month old mice, with *h*_α_*-Syn*^*+*^*:Nrf2*^*-/-*^ mice having significantly higher mRNA levels than wildtype (**Figure 1A-B**). Interestingly 4-hydroxynonenal (4-HNE) adduct formation was similar across all groups in the MB, but was higher in the ST of 3 month old *h*_α_*-Syn*^*+*^*:Nrf2*^*-/-*^ than *h*_α_*-Syn*^*+*^*:Nrf2*^*+/+*^ mice, which was further increased in both genotypes at 6 months (**Figure 1A**). In addition, Aifm2/Fsp1 (recently identified to be an anti-ferroptotic lipophilic antioxidant) protein levels were significantly higher in the MB and ST of 3 month old *h*_α_*-Syn*^*+*^*:Nrf2*^*-/-*^ mice, as well as both genotypes at 6 months of age (**Figure 1A**). Please note, the mRNA and protein levels of the key anti-ferroptotic mediator Gpx4 were similar across all genotypes and brain regions at both timepoints (**Figure 1A-B**).

**Figure 1:**
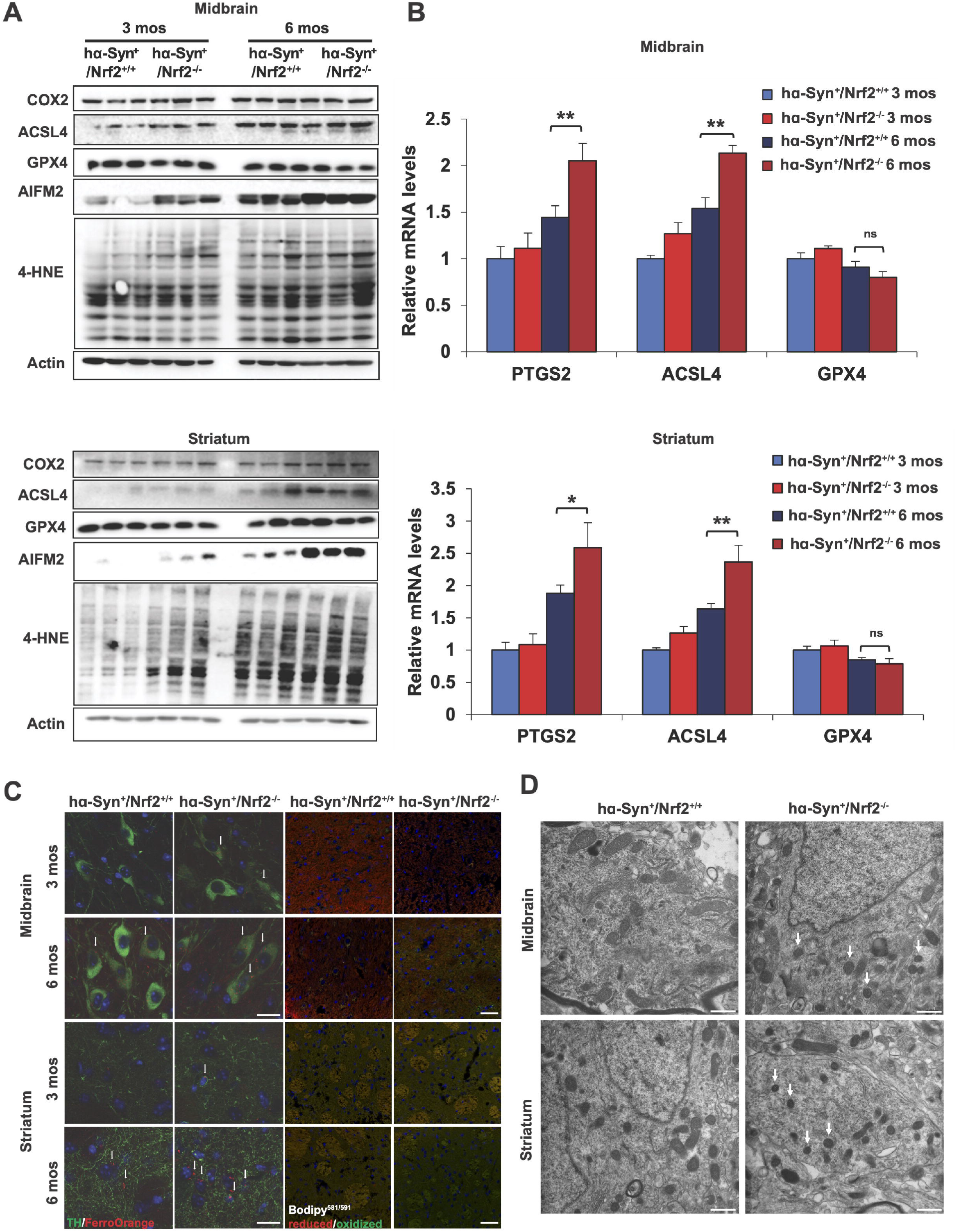
Loss of Nrf2 increases markers of ferroptosis in the PD brain. The midbrain (MB) and striatum (ST) of 3 and 6 month old *h*α*-Syn*^*+*^*:Nrf2*^*+/+*^ and *h*α*-Syn*^*+*^*:Nrf2*^*-/-*^ mice were collected and assessed for key markers of ferroptosis. (A) Immunoblot analysis of Cox2, Acsl4, Gpx4, xCT, Aifm2, and 4-HNE adduct protein levels. β-actin was used as an internal loading control. (B) qRT-PCR of *Ptgs2, Acsl4*, and *Gpx4* mRNA levels. n=5 (C) Immunofluorescent staining of tyrosine hydroxylase (green) and FerroOrange (orange; left panels), or C11-BODIPY^581/591^ (red = reduced, green = oxidized; right panels). Scale bar = 50 µm. (D) TEM micrographs of the MB and ST of 6-month-old *h*_α_*-Syn*^*+*^*:Nrf2*^*+/+*^ and *h*_α_*-Syn*^*+*^*:Nrf2*^*-/-*^ mice. Arrows indicate condensed mitochondria. Scale bar = 500 nm. *p<0.05, **p<0.01, One-way ANOVA with Tukey’s post-hoc test.

As increased free iron and lipid peroxidation are key indicators of ferroptosis, ferrous (Fe^2+^) iron and lipid peroxide levels were measured via FerroOrange and Bodipy^581/591^, respectively. In the MB and ST, FerroOrange levels were significantly increased in the *h*_α_*-Syn*^*+*^*:Nrf2*^*-/-*^ mice, particularly at 6 months of age (**Figure 1C**). Importantly, the increased FerroOrange signal observed in the MB colocalized with tyrosine hydroxylase-positive neurons, indicating that ferroptosis is occurring in dopaminergic neurons, a key cell type affected during PD progression. A similar increase in Bodipy^581/591^ levels in the MB and ST was observed in both the 3 month and 6 month old *h*_α_*-Syn*^*+*^*:Nrf2*^*-/-*^ mice, with the most significant change occurring in the ST (**Figure 1C**). The observed increase in free iron may be the result, at least in part, of ferritinophagy blockage, as LC3-II (indicator of autophagosomes), Ncoa4 (adaptor for ferritin removal via autophagy), and Fth1 (ferritin heavy chain) protein levels were all higher in both brain regions of 6 month *h*_α_*-Syn*^*+*^*:Nrf2*^*-/-*^ animals (**Figure S1A**). Finally, mitochondrial morphology in the MB and ST of 6-month-old animals was assessed via transmission electron microscopy, as shrunken mitochondria with increased membrane density has been reported to be associated with ferroptotic death [6]. Mitochondria in both the MB and ST of the *h*_α_*-Syn*^*+*^*:Nrf2*^*-/-*^ mice were significantly smaller and more dense than the *h*_α_*-Syn*^*+*^*:Nrf2*^*+/+*^ mice (**Figure 1D**). Mitochondrial morphology was also significantly altered in the hippocampus and cortex (**Figure S1B**). Altogether, these data indicate that excess _α_-syn promotes ferroptosis in an age- and NRF2-dependent manner.

### h_α_-Syn^+^:Nrf2^-/-^ mice exhibit glutathione depletion and increased oxidative stress at 6 months of age

Next, we determined if loss of NRF2 was increasing ferroptotic markers due to decreased target gene expression and subsequent accumulation of reactive species. The protein levels of both Akr1c1 and Gclm, two well-established NRF2 target genes, were both decreased in the 3 and 6 month old *h*_α_*-Syn*^*+*^*:Nrf2*^*-/-*^ mice compared to the *h*_α_*-Syn*^*+*^*:Nrf2*^*+/+*^ mice (**Figure 2A**). Interestingly, a decrease in the mRNA level of *Akr1c1* and *Gclm* was only observed in the 6 month animals, when *h*_α_*-Syn*^*+*^*:Nrf2*^*-/-*^ was compared to *h*_α_*-Syn*^*+*^*:Nrf2*^*+/+*^ mice (**Figure 2B**). Manganese superoxide dismutase (*Sod2*/MnSOD) levels, an important mitochondrial antioxidant enzyme and potential NRF2 target, were also significantly decreased at the mRNA level in the 6 month old *h*α*-Syn*^*+*^*:Nrf2*^*-/-*^ mice (**Figure 2B**). As expected based on diminished expression of protective target genes, reactive oxygen species (ROS), malondialdehyde (MDA) formation, and free iron were all significantly increased in the MB and ST of 6 month old *h*_α_*-Syn*^*+*^*:Nrf2*^*-/-*^ mice, which corresponded with a decrease in total glutathione (GSH) levels (**Figure 2C-F**). Similar changes in these parameters were also observed in the hippocampus and cortex (**Figure S2A-D**). Thus, loss of NRF2 significantly increases the iron, ROS, and lipid peroxides that promotes ferroptosis.

**Figure 2:**
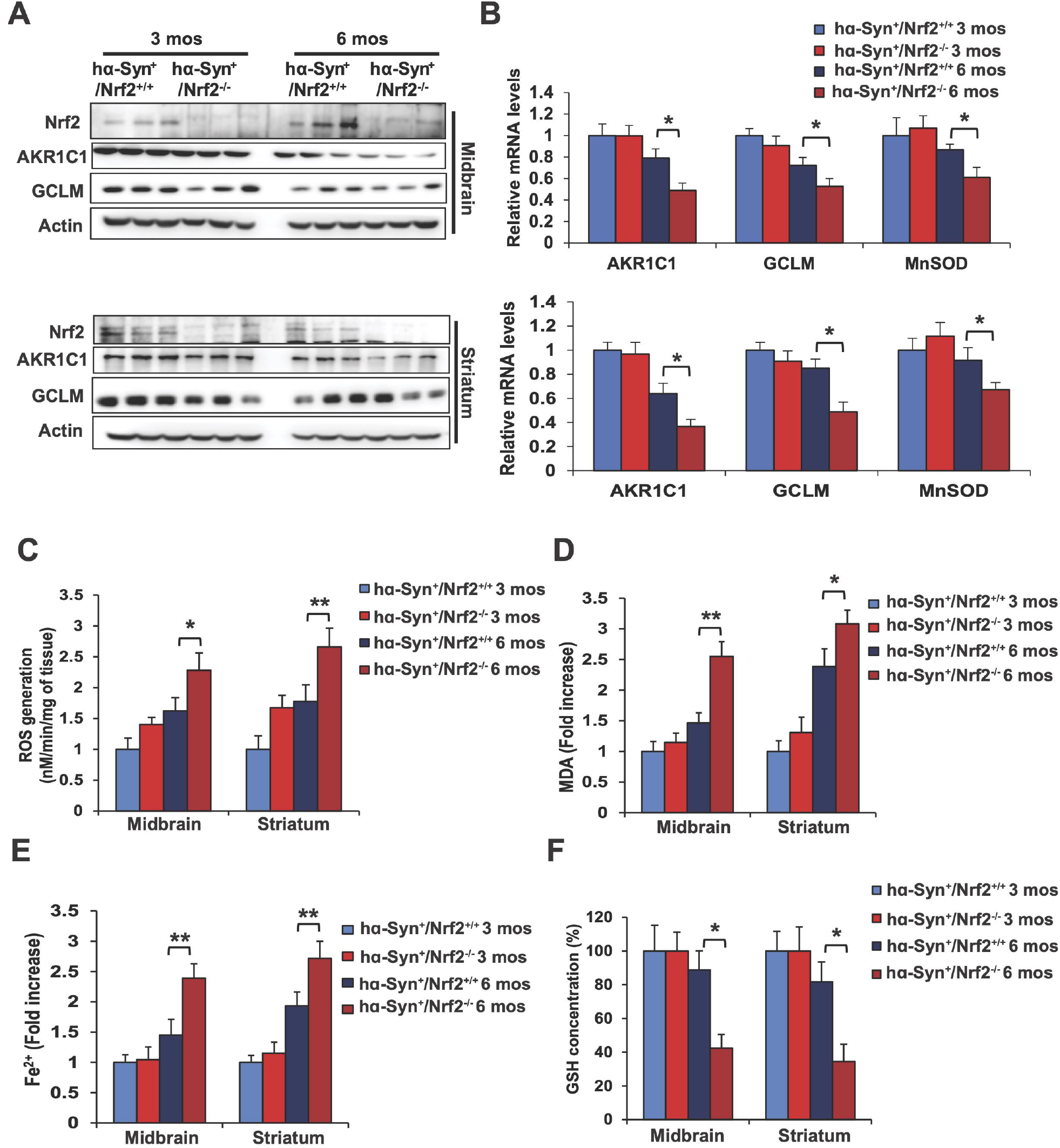
6-month-old *h*α*-Syn*^*+*^*:Nrf2*^*-/-*^ mice exhibit increased free iron, ROS, and lipid peroxidation. The midbrain (MB) and striatum (ST) of 3 and 6 month old *h*_α_*-Syn*^*+*^*:Nrf2*^*+/+*^ and *h*_α_*-Syn*^*+*^*:Nrf2*^*-/-*^ mice were collected and assessed for changes related to loss of NRF2. (A) Immunoblot analysis of Nrf2, and two of its known target genes, Akr1c1 and Gclm. β-actin was used as an internal loading control. (B) qRT-PCR analysis of *Akr1c1, Gclm*, and *Sod2* (MnSOD). n=5 (C) ROS generation measured by electron paramagnetic resonance (EPR) spectroscopy. n=6 (D) Malondialdehyde (MDA) formation measured by TBARs assay. n=6 (E) Free ferrous iron (Fe^2+^) measured via Ferene-S based colorimetry. n=6 (F) Total glutathione (GSH) levels measured via Quantichrom assay. n=6 *p<0.05, **p<0.01, One-way ANOVA with Tukey’s post-hoc test.

### h_α_-Syn^+^:Nrf2^-/-^ mice exhibit an age-dependent exacerbation of α-syn-induced pathogenesis

We have shown previously that *h*_α_*-Syn*^*+*^*:Nrf2*^*-/-*^ mice at 3 months old have greater α*-*syn associated pathology and behavioral deficits than their wildtype counterparts [9]. To determine if the observed age-dependent increase in enhanced ferroptosis markers was associated with more severe α*-*syn pathology, α*-*syn phosphorylation and accumulation, as well as behavioral deficits were assessed. Similar to our previous reports, 3-month-old *h*_α_*-Syn*^*+*^*:Nrf2*^*-/-*^ mice had higher phosphorylated _α_*-*syn levels, as well as increased formation of oligomeric α*-*syn species in the MB, but not ST (**Figure 3A**). 3-nitrotyrosine levels (3-NT), a PD-relevant indicator of oxidative damage, was also increased in the 3-month MB, but not ST, compared to wildtype (**Figure 3A**). At the 6 month time point, phosphorylated and oligomeric _α_*-*syn, as well as 3-NT levels, were all significantly higher in both the MB and ST than at 3 months, with the *h*_α_*-Syn*^*+*^*:Nrf2*^*-/-*^ mice exhibiting the most severe phenotype (**Figure 3A**). Similarly, FerroOrange fluorescence was more pronounced in both genotypes at 6 months of age, and colocalized with phospho-α*-*syn, indicating that iron is accumulating in cells where pathogenic alterations to α*-*syn are occurring (**Figure 3B**). Phenotypically, nestlet pulldown/nest building, errors per step while traversing a beam, and number of entries/time spent in the center of an open field were all significantly worse in 6 month old *h*_α_*-Syn*^*+*^*:Nrf2*^*-/-*^ mice than wildtype, whereas beam traverse time and time spent grooming showed no significant difference (**Figure 3C-E**). These results indicate that loss of NRF2 increases α*-*syn pathology and induction of behavioral deficits, with older mice showing a more pronounced phenotype.

**Figure 3:**
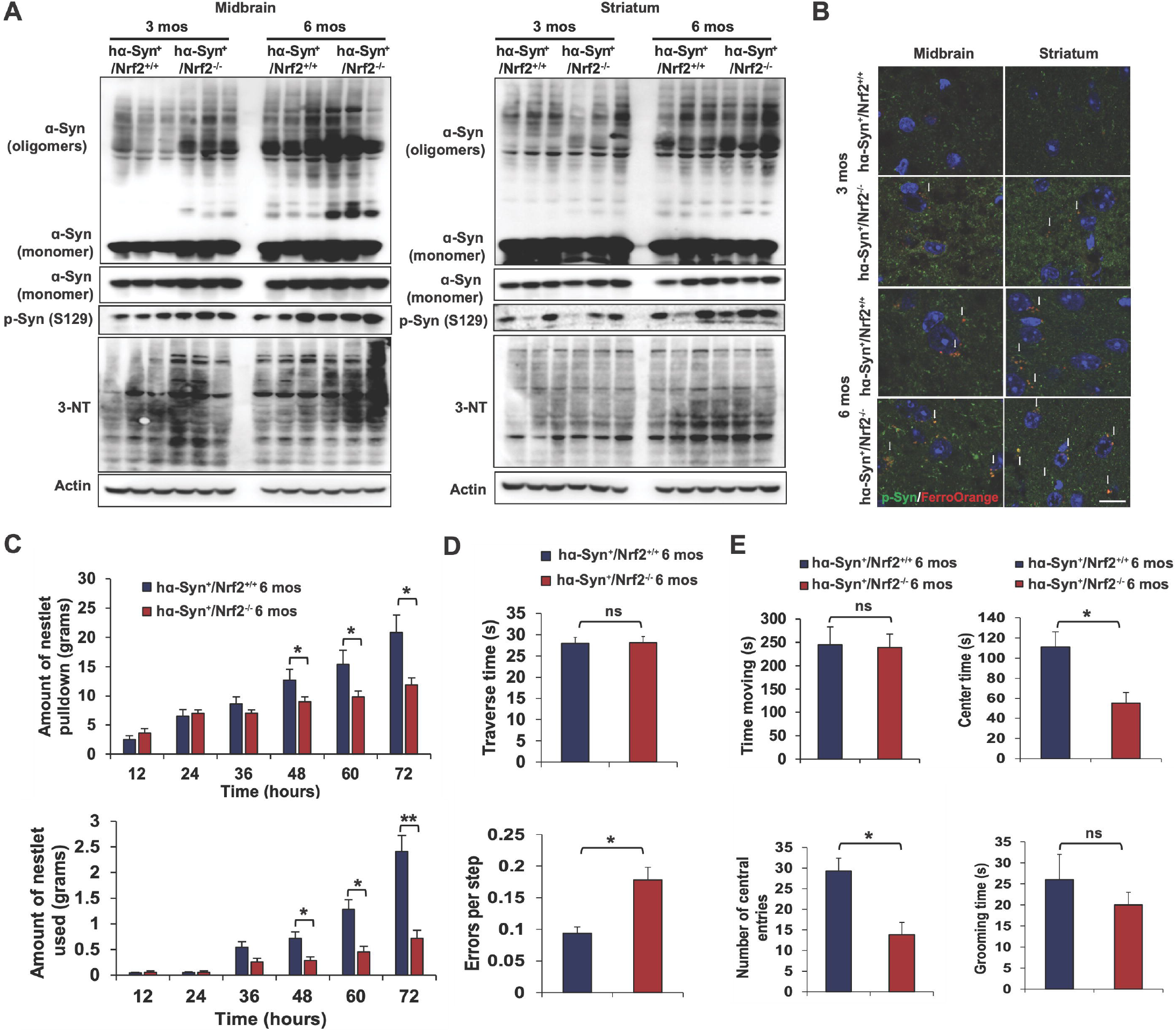
6-month-old *h*α*-Syn*^*+*^*:Nrf2*^*-/-*^ mice exhibit greater □-syn pathology and behavioral deficits. The midbrain (MB) and striatum (ST) of 3 and 6 month old *h*_α_*-Syn*^*+*^*:Nrf2*^*+/+*^ and *h*_α_*-Syn*^*+*^*:Nrf2*^*-/-*^ mice were collected and assessed for changes in L-syn accumulation/phosphorylation and indicators of pro-ferroptotic stress. (A) Immunoblot analysis of monomeric and oligomeric □-syn, phosphorylated □-syn (S129), and 3-nitrotyrosine (3-NT). β-actin was used as an internal loading control. (B) Immunofluorescent staining of phosphorylated □-syn (green) and FerroOrange (orange). Scale bar = 50 µm. 6 month old *h*_α_*-Syn*^*+*^*:Nrf2*^*+/+*^ (n=16) and *h*_α_*-Syn*^*+*^*:Nrf2*^*-/-*^ (n=16) mice were subjected to a battery of behavioral tests. (C) Nestlet pulled down (top panel) and nestlet used (bottom panel). (D) Narrow beam traverse time (top panel) and foot slips/errors per step (bottom panel). (E) Time spent moving (top left panel), center time (top right panel), and number of central entries (bottom left panel) during an open field test, and grooming time (bottom right panel). For (C), *p<0.05, **p<0.01 Two-way repeated measure ANOVA. For (D-E), *p<0.05, Unpaired Student’s *t*-test.

### Loss of NRF2 enhances susceptibility to ferroptosis induction ex vivo

Next, the effect of genetic ablation of NRF2 and α*-*syn overexpression on enhancing susceptibility to the well-established ferroptosis inducer erastin was determined using *ex vivo* brain slices. Intriguingly, *h*_α_*-Syn*^*+*^*:Nrf2*^*+/+*^ brain slices exhibited more significant erastin-induced ferroptotic death than their *h*α*-Syn*^*-*^*:Nrf2*^*+/+*^ counterparts, as indicated by an increase in the number of propidium iodide-positive cells and decrease in NeuN signal (neuronal marker) following treatment (**Figure 4A-B**). The *h*α*-Syn*^*+*^*:Nrf2*^*-/-*^ and *h*α*-Syn*^*-*^*:Nrf2*^*-/-*^ brain slices both displayed higher levels of ferroptotic death than wildtype, with the anti-ferroptotic iron chelator deferoxamine (DFO) reversing cell death across all groups (**Figure 4A-B**). The observed changes in erastin-induced ferroptotic death were associated with a similar change in *Ptgs2* and *Acsl4* mRNA levels, with the *Syn*^*+*^ groups trending higher than the *Syn*^*-*^ groups, and *h*_α_*-Syn*^*+*^*:Nrf2*^*-/-*^ brain slices exhibited the highest levels of these pro-ferroptotic markers (**Figure 4C**). This phenotype was also validated at the protein level, as Cox2 and Acsl4 protein levels followed the same trend as the mRNA, and 4-HNE protein adduct formation was also significantly higher in the *Syn*^*+*^*:Nrf2*^*-/-*^ groups, and decreased upon cotreatment with DFO (**Figure 4D**). This indicates that both loss of NRF2 and _α_*-*syn overexpression can significantly induce neuronal ferroptotic cell death in an *ex vivo* setting.

**Figure 4:**
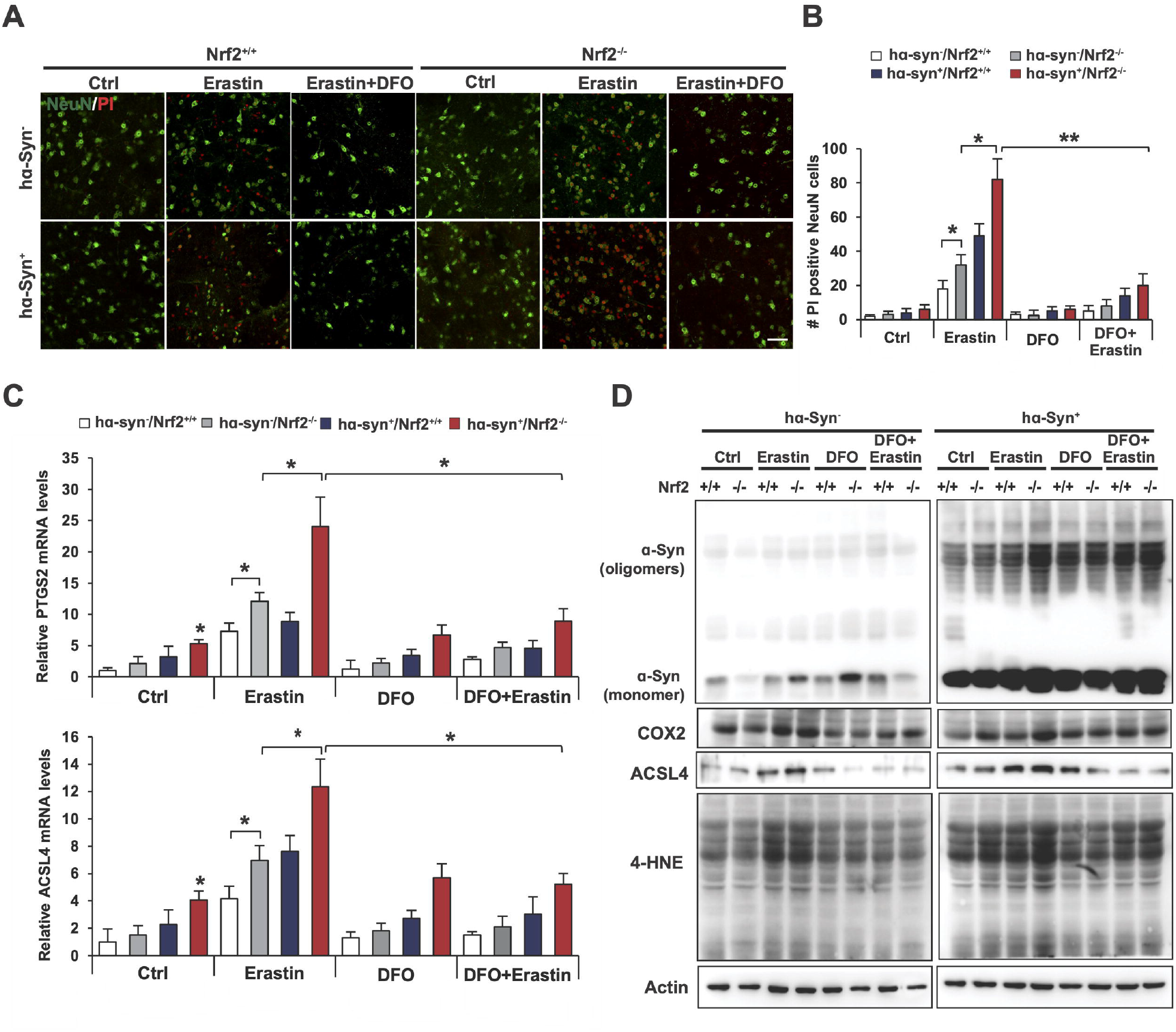
Loss of Nrf2 increases susceptibility to erastin-induced ferroptosis *ex vivo*. Brain slices were prepared from 6 month old *h*_α_*-Syn*^*-*^*:Nrf2*^*+/+*^, *h*_α_*-Syn*^*-*^*:Nrf2*^*-/-*^, *h*_α_*-Syn*^*+*^*:Nrf2*^*+/+*^, and *h*_α_*-Syn*^*+*^*:Nrf2*^*-/-*^ mice and were either left untreated or treated with10 µM deferoxamine (DFO) for 1 hr, followed by treatment with 10 µM erastin for 16 hr. (A) Immunofluorescent staining of NeuN (green; neuronal marker) and propidium iodide (PI, red; marker of cells undergoing ferroptosis). (B) qRT-PCR of *Ptgs2* (top panel) and *Acsl4* (bottom panel) mRNA levels. n=3 (C) Immunoblot analysis of monomeric and oligomeric □-syn, Cox2, Acsl4, and 4-HNE adduct protein levels. β-actin was used as an internal loading control. *p<0.05, **p<0.01, Two-way ANOVA Holm-Sidak post hoc test.

### _α_-syn enhances ferroptosis and suppresses NRF2 protein levels in neuronal cells

To further validate the NRF2-α*-*syn-ferroptosis axis, differentiated SH-SY5Y neuroblastoma cells were transduced with adenovirus encoding either WT or A53T (higher aggregation potential) α*-*syn, and indicators of ferroptosis following treatment with known ferroptosis inducers were assessed. Similar to our *in vivo* and *ex vivo* studies, overexpression of both forms of α*-*syn resulted in a significant increase in the number of ferroptotic cells following treatment with RSL-3, erastin, or cystine-depletion (CD) (**Figure 5A-B**). Furthermore, α*-*syn-enhanced ferroptosis could be reversed by treatment with DFO, or the lipophilic antioxidant ferrostatin-1 (Fer-1) (**Figure 5C**). Next, two different *in vitro* models of α*-*syn overexpression were used to check α*-*syn effects on NRF2 function. Surprisingly, both transduction with adenovirus encoding WT α*-*syn, as well as doxycycline induction of α*-*syn expression, resulted in a significant decrease in the protein levels of NRF2 and its target genes AKR1C1 and GCLM (**Figure 5D-E**). Intriguingly, the decrease in NRF2 protein level was independent of changes in KEAP1 protein level, or *NFE2L2*/NRF2 mRNA levels (**Figure 5E-F**). Finally, the mRNA levels of *PTGS2* and *ACSL4* were increased, whereas *AKR1C1* and *GCLM*, but not *NQO1*, were all decreased (**Figure 5F**). GPX4 levels remained unchanged, except for a slight decrease at the mRNA level at the 72 hr induction time point. This indicates that α*-*syn accumulation can suppress NRF2 protein levels, decreasing target gene expression, and enhancing susceptibility to ferroptotic cell death.

**Figure 5:**
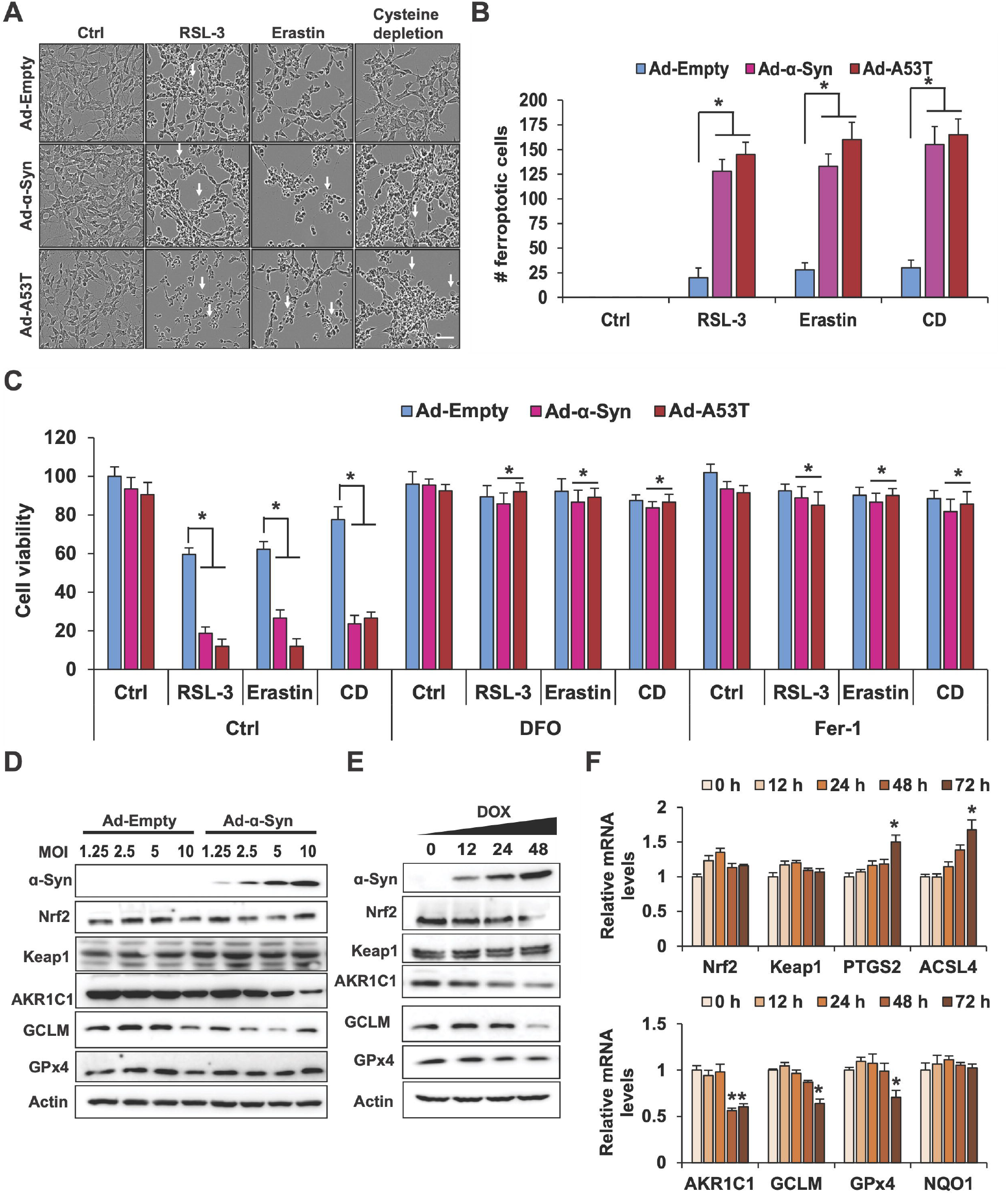
□-syn overexpression enhances susceptibility to ferroptosis and suppresses NRF2 in a neuronal cell type. SH-SY5Y neuroblastoma cells were differentiated for 7 days, transduced with adenovirus (Ad) encoding WT or A53T L-syn for 24 hr, then treated with 0.5 µM RSL-3, 1 µM erastin, or cystine depletion (CD) for 24 hr. (A) Representative images of ferroptotic cell morphology (ballooning) taken using the Incucyte Bioimaging system. Scale bar = 100 µm. (B) Number of ferroptotic cells from (A). (C) Cells were treated the same as (A-B) in the presence or absence of 100 µM DFO or 10 µM ferrostatin-1 (Fer-1) and cell viability was measured via MTT assay. n=3 (D) Immunoblot analysis of monomeric □**-**syn, NRF2, KEAP1, AKR1C1, xCT, GLCM, and GPX4 protein levels from SH-SY5Y cells transduced with increasing MOIs of adenovirus encoding either empty vector or WT □-syn for 24 hr. Doxycycline-inducible □**-**syn SH-SY5Y cells were induced with doxycycline for the indicated time points. (E) Immunoblot analysis of monomeric □**-**syn, NRF2, KEAP1, AKR1C1, xCT, GLCM, and GPX4 protein levels. (F) qRT-PCR of *NFE2L2/*NRF2, *KEAP1, PTGS2, ACSL4, AKR1C1, GCLM, GPX4*, and *NQO1*. (B and D) *p<0.05, **p<0.01, One-way ANOVA with Tukey’s post-hoc test. (C) *p<0.05, Two-way ANOVA Holm-Sidak post hoc test.

## Discussion

In this study, we identified NRF2 as a key mediator of neuronal ferroptosis, particularly in a PD-relevant context where excess α*-*syn is present. Using *in vitro, ex vivo*, and *in vivo* models of PD, we show that there is a vicious cycle of α-syn overexpression, NRF2 suppression, and enhanced neuronal ferroptotic cell death that promotes the onset of parkinsonian phenotypes. To our knowledge, this is the first time that the following has been demonstrated. Firstly, that loss of NRF2 can drive α*-*syn-promoted ferroptosis, and secondly, that α*-*syn accumulation can suppress NRF2 function. The latter is perhaps one of the most striking findings of this study, particularly considering that increased α*-*syn can suppress NRF2 in the absence of any changes to its mRNA levels or KEAP1 protein expression. While beyond the scope of the current study, determining if there is a physical interaction between NRF2 and α*-*syn that affects its stability, or if some other mechanism of KEAP1-independent degradation is occurring, will be interesting to pursue. Regardless, the important findings from this study are that loss of NRF2 enhances neuronal susceptibility to ferroptotic death, and that α*-*syn suppression of NRF2 function could be a key driver of age-related PD pathogenesis.

The discovery that α*-*syn can promote ferroptosis *in vivo*, as well as a clear indication that an age-dependent decline in NRF2 could enhance neuronal death by this mechanism, is important for several reasons. First and foremost, a critical limitation in treating PD is that by the time many of the hallmark symptoms are diagnosed, irreversible neuronal loss and damage to critical brain regions has already occurred. However, despite decades of knowing that neuronal cell death is a critical driver of PD progression, targetable mechanisms to prevent neuronal loss remain limited. Another important aspect of this study is the indication that NRF2 is critical to prevent neuronal ferroptosis, and NRF2 induction has long been proposed as a therapeutic approach for PD. Regrettably, one longstanding problem with targeting NRF2 in a neurodegenerative disease context is a lack of compounds that can successfully pass the blood brain barrier (BBB). Thus, another important need is not only the discovery of safe inducers that can cross the BBB, but also determining ways to target the downstream pathways affected by loss of NRF2, so that alternative therapies that mitigate disease progression might be discovered. Thus, the NRF2-α*-*syn-ferroptosis axis represents a viable target for therapeutic generation, as both NRF2 induction or pharmacological inhibition of ferroptosis, could avert neuronal death and delay or even prevent PD onset and progression.

## Materials and Methods

### Chemicals and Reagents

Erastin, ferrostatin-1 (Fer-1), and 3-(4,5-Dimethylthiazol-2-yl)-2,5-diphenyltetrazolium bromide (MTT) were purchased from Sigma. 1-hydroxy-3-methoxycarbonyl-2,2,5,5-tetramethylpyrrolidine (CMH; NOX-0.2), deferoxamine (DFO), diethyldithiocarbamic acid (DETC), and Krebs-HEPES buffer (NOX-07.6.1) were purchased from Noxygen.

### Cell culture

Human dopaminergic neuroblastoma cells (SH-SY5Y) were obtained from the American Type Culture Collection (ATCC; Manassas, VA). Cells were cultured in DMEM/F12 medium containing 10% fetal bovine serum, 100 units/ml penicillin-streptomycin in a humidified 37°C incubator with 5% CO2. The dox-(inducible) α-synuclein cells were generated as describe previously [21], and were kindly provided by Dr. Muralidhar L. Hegde. For the differentiated SH-SY5Y experiments, cells were treated with 10 μM retinoic acid (RA) for seven days to induce neuronal differentiation.

### Recombinant Adenoviral vectors

Replication-deficient recombinant adenoviruses (pAd5CMV) encoding wild type (WT) or mutant A53T α-synuclein were generated as described previously [22], and were a kind gift from Dr. Jean-Christophe Rochet (Purdue University). Adenoviruses were amplified and tittered as described previously [23]. Cells were transduced with adenoviral vectors at the indicated multiplicity of infection (MOI) for 24 h, washed, and subsequently treated under the specified experimental conditions.

### Animals

Hemizygous human Thy1-α-Syn mice (hα-Syn+; C57Bl6/DBA background; founder breeder pairs were originally provided by Dr Marie-Francoise Chesselet, UCLA) [24]. Nrf2 knockout (*h*_α_*-Syn*^*+*^;*Nrf2*^*-/-*^) and wild-type (*h*_α_*-Syn*^*+*^;*Nrf2*^*+/+*^) mice were generated as described previously [9]. Mice were handled according to the rules and regulations of the NIH and Institutional Guidelines on the Care and Use of Laboratory Animals. All experimental protocols were approved by the University of Arizona Institutional Animal Care and Use Committee. The mice were housed under a reverse 12-hour light-dark cycle condition with food and water available ad libitum. Male littermates were used in the study, and the genotypes of all mice were verified with PCR analysis of tail DNA.

### Experimental Design

All four genotypes of mice produced were maintained until 3 and 6 mos of age, at which point they were subjected to behavioral tests. Subsequently, animals were sacrificed at 3.5 and ∼6.5 mos of age and the indicated brain regions were processed for histological and molecular analysis. For the histological studies, mice were sacrificed using sodium pentobarbital (60 mg/kg) and perfusion with 4% paraformaldehyde (PFA). Subsequently, brains were extracted and post-fixed in 4% PFA, sunk through a 30% sucrose solution, and sectioned (30 μm) in the coronal plane on a freezing sliding microtome. For molecular studies, animals were sacrificed using sodium pentobarbital (60 mg/kg), brains extracted and subjected to microdissection on ice to isolate tissues from the midbrain, striatum, cortex, and hippocampus. The dissected tissues were then snap frozen in liquid nitrogen for later molecular analyses.

### Behavioral analyses

#### Nest Building Task

Nest building is a natural motor behavior requiring the use of orofacial and forelimb movements that can be used to assess PD relevant sensorimotor function in rodents [25]. This task tests the ability of the animal to retrieve and use woven cotton pads (nestlets) placed in their cage’s feeding bin. Once the nestlets are grasped and placed into the cage, the animals pull the nesting material apart with their forelimbs and teeth, breakdown the cotton in their mouths, and incorporate it into their bedding. Thus, forelimb dexterity and fine motor behavior can be measured with this task. Briefly, mice are transferred to individual testing cages approximately 1 hr before the dark phase and weighed nestlets are placed in the feeder bin of the cage (5 nestlets, ∼12g/cage). The amount of nestlets that were retrieved (nestlet pulldown) and broken down (nestlet usage) for nest building was measured every 12 hours over a 72 hour period (12, 24, 36, 48, 60 and 72 hours) by weighing the unused material in the feeder and inside the cage as reported previously [9]. Data were presented as the amount of nestlet pulled down and the amount used.

#### Challenging Beam Task

Mice were subjected to a challenging beam task to test motor coordination and agility [25]. A wooden beam consisting of four continuous sections (25 cm in length each, 1m total length), which incrementally narrows from a starting width of 3.5 cm to an ending width of 0.5 cm, was used. The widest segment of the beam served as a loading platform for the animals and the narrowest end connected into the home cage. Mice received 2 days of training before testing. During the training phase, animals were brought to the behavior room at the start of their dark cycle and acclimated to the testing room for ∼2 hrs each day. The animal was placed on the beam at its widest point and trained to move across the beam towards the narrowest part and into the home cage. Testing started on day 3 when mice underwent 5 unassisted trials on the beam. To increase difficulty, a mesh grid (1 cm2 grid) of corresponding width was placed over the beam surface leaving an ∼1 cm space between the grid and the beam surface to allow adequate visualization of the mice’s paws as they walked. Each trial was videotaped, then viewed and rated in slow motion for foot slip errors, number of steps made by each animal to cross the beam, and time taken to traverse the beam, by an investigator blind to the mouse genotype. A foot slip error was counted when, during a forward movement, a limb (forelimb or hindlimb) slid through the grid and was visible between the grid and the beam surface. By scoring each limb slip individually, the severity of the error could be measured. The trial ended when an animal fell off the beam, reached the maximum allowed time (60 sec), or traversed the full distance. Errors per step, time to traverse, and number of steps were calculated as the average across all five trials.

#### Open field task

The open field test was used to determine spontaneous exploratory activity of the mice [26]. Activity was assessed in an open arena (60 wide x 60 long x 60 cm high) with a central box drawn in the middle of the field floor (30 × 30 cm center). Animals were tested within the first 2-4 h of the dark cycle after being habituated to the testing room for 15 min. Mice were placed individually in the center of the open field (30 cm square) and their movement monitored (videotaped) for 15 min. Videos were analyzed for the following parameters: time spent moving, distance traveled, number of times the central box is encroached (center entries), time spent in central box (center time), number of rears, and time spent grooming.

### Cell viability and ferroptotic cell count

Cell viability was analyzed using an MTT assay. Briefly, cells were seeded in a 96 well plate. 24 h after treatment, 20 µL of MTT (5 mg/mL) in 1X PBS was added directly to the cell culture media and allowed to incubate for 3 h. Media was then removed, and solvent (isopropanol/HCl) was added to the cells and absorbance was measured at 570 nm via a SpectraMax iD5 MultiMode Microplate Reader (Molecular Devices). Ferroptotic cell death was monitored continuously using the IncuCyte (Essen Biosciences). The number of ferroptotic cells was counted manually based on ferroptotic cell morphology (ballooning).

### Glutathione (GSH) Assay

Intracellular glutathione concentrations were measured using the QuantiChrom glutathione assay kit from BioAssay Systems. The procedure was performed according to the manufacturer’s instructions. Briefly, 5,5’-dithiobis 2-nitro benzoic acid (DTNB) reacts with intracellular GSH to form a yellow product. Absorbance was measured at 412 nm and GSH in cellular extracts was quantified via comparison to a calibration curve generated using GSH as a standard. Data were normalized to tissue weight.

### Lipid Peroxidation Assay

The relative intracellular malondialdehyde (MDA) concentration was assessed using a Lipid Peroxidation (MDA) Assay Kit from Sigma-Aldrich. Briefly, MDA in the sample reacts with thiobarbituric acid (TBA) to generate an MDA-TBA adduct. The MDA-TBA adduct can be quantified colorimetrically (OD=532 nm). Lipid Peroxidation was also measured by C11-BODIPY^581/591^ dye (Thermo Fisher Scientific). Briefly, sections were blocked [10% normal goat serum, 0.5% Triton-X-100 in Tris buffered saline (TBS, pH 7.4)] and incubated in 20 µM C11-BODIPY^581/591^ for 30 mins. Oxidation of the polyunsaturated butadienyl portion of the dye results in a shift of the fluorescence emission peak from _∼_590 nm to _∼_510 nm.

### Iron assay

Intracellular ferrous (Fe^2+^) iron levels were determined by colorimetric assay using Ferene-S (Sigma-Aldrich). In this assay, iron is released by the addition of an acidic buffer. Ferrous iron is reacted with a chromagen resulting in a colorimetric (593 nm) product, proportional to the iron present. In another assay, intracellular chelatable iron was determined using the fluorescent probe FerroOrange from DOJINDO. Briefly, sections were blocked [10% normal goat serum, 0.5% Triton-X-100 in Tris buffered saline (TBS, pH 7.4)] and incubated in 1 µM ferroOrange for 30 mins. FerroOrange is a fluorescent probe that enables cell fluorescent imaging of intracellular Fe^2+^ (ex: 543 nm, em: 580 nm).

### Electron Paramagnetic Resonance (EPR)

EPR was performed as described previously [9]. Briefly, brain tissues were dissected (midbrain, striatum, hippocampus, and cortex), and incubated with spin trap in the presence of metal chelators (200 µM cyclic hydroxylamine 1-hydroxy-. 3-methoxycarbonyl-2,2,5,5-tetramethylpyrrolidine [CMH], 25 µM deferoxamine [DFO], and 5 µM diethyldithiocarbamate [DETC] in filtered 20 mM KREBS-HEPES buffer) (Noxygen) for 30 min prior to measurement. Buffer was then collected, and changes in CMH oxidation were measured for 15 min using the e-scanM Multipurpose Bench-top EPR system (Noxygen Science and Transfer Diagnostics GmbH). The relative production of reactive oxygen species was represented as the nanomolar concentration of oxidized spin trap divided by the time of trap incubation normalized to the total milligrams of tissue, then data were normalized to control.

### Immunohistochemistry

Immunohistochemistry was performed using previously established methods [9, 27]. Briefly, sections were blocked [10% normal goat serum, 0.5% Triton-X-100 in Tris buffered saline (TBS, pH 7.4)] and incubated in primary antibodies anti-phospho α-Syn S129 (1:300 - ab-59264, Abcam, Cambridge, UK), and anti-tyrosine hydroxylase (1:4000 - MAB318, Chemicon Temecula, CA) overnight at room temperature (RT). Primary antibodies were detected in a 2 hr incubation at RT with secondary antibodies coupled to fluorochromes Alexa 488 or 555 (Life Technologies-Molecular Probes, Grand Island, NY) and counterstained with 4’,6’-diamidino-2-phenylindole, dihydrochloride (DAPI, Life Technologies). For imaging, a Leica SP5-II confocal microscope (Leica Microsystems, Wetzlar, Germany) at 20X or 60X magnification was used, and Z sectioning was performed at 1-2 μm intervals in order to verify the co-localization of markers. Image extraction and analysis was conducted via the Leica LAS software (LAS AF 2.7.3.9723).

### Real-Time qRT-PCR

Total mRNA was extracted using TRIzol (Invitrogen, Carlsbad, CA) according to the manufacturer’s instructions. cDNA was then synthesized using 2 μg of mRNA and a Transcriptase first-strand cDNA synthesis kit (Promega, Madison, WI). Real-Time qPCR to detect the indicated targets was performed on a LightCycler® 480 System (Roche Life Science, Penzberg, Germany) using PowerUp SYBR Green Master Mix (Thermo Fisher Scientific, Waltham, MA). Actin or GAPDH were used as an internal control. The reaction conditions were as follows; UDG activation 50°C (2 min), Dual-Lock™ DNA polymerase 95 °C (2 min), Denaturation 95 °C (15 sec), Annealing 55-60°C (15 sec), Extension 72°C (1 min) and 40 cycles. All experiments were performed in triplicate. Relative expression levels were calculated using the 2-ΔΔCT method. Primer sequences (5′-3′) were as follows: mouse-*Ptgs2*-Forward 5’-CATCCCCTTCCTGCGAAGTT-3’ mouse-*Ptgs2*-Reverse 5’-CATGGGAGTTGGGCAGTCAT-3’ mouse-*Acsl4*-Forward 5’-GGAATGACAGGCCAGTGTGA-3’ mouse-*Acsl4*-Reverse 5’-GAGGGGCGTCATAGCCTTTC-3’ mouse-*Gpx4*-Forward 5’-GGTTTCGTGTGCATCGTCAC-3’ mouse-*Gpx4*-Reverse 5’-GGGCATCGTCCCCATTTACA-3’ mouse-*Slc7a11*-Forward 5’-TGCAATCAAGCTCGTGAC-3’ mouse-*Slc7a11*-Reverse 5’-AGCTGTATAACTCCAGGGACTA-3’ mouse-*Akr1c1*-Forward 5’-CCGGTATGCAACCAGGTAGA-3’ mouse-*Akr1c1*-Reverse 5’-AGACTGGTGTTGAGGACCAC-3’ mouse-*Gclm*-Forward 5’-ATGGAGGCGATGTTCTTGAG-3’ mouse-*Gclm*-Reverse 5’-GTCTCCAGAGGGTCGGAT-3’ mouse-*MnSOD*-Forward 5’-GCGGTCGTGTAAACCTCAAT-3’ mouse-*MnSOD*-Reverse 5’-GCCATAGTCGTAAGGCAGGT-3’ mouse-*Actin*-Forward 5’-AAGGCCAACCGT GAAAAGAT-3’ mouse-*Actin*-Reverse 5’-GTGGTACG ACCAGAGGCATAC-3’ human-*NRF2*-Forward 5’-ACACGGTCCACAGCTCATC -3’ human-*NRF2*-Reverse 5’-TGTCAATCAAATCCATGTCCTG -3’ human-*KEAP1*-Forward 5’-ACCACAACAGTGTGGAGAGGT -3’ human-*KEAP1*-Reverse 5’-CGATCCTTCGTGTCAGCAT -3’ human-*ACSL4*-Forward 5’-GGAATGACAGGCCAGTGTGA-3’ human-*ACSL4*-Reverse 5’-TCACCAGTGCAAAACCACCT-3’ human-*PTGS2*-Forward 5’-GTTCCACCCGCAGTACAGAA-3’ human-*PTGS2*-Reverse 5’-AGGGCTTCAGCATAAAGCGT-3’ human-*AKR1C1*-Forward 5’-CATGCCTGTCCTGGGATTT-3’ human-*AKR1C1*-Reverse 5’-AGAATCAATATGGCGGAAGC-3’ human-*GCLM*-Forward 5’-GACAAAACACAGTTGGAACAGC-3’ human-*GCLM*-Reverse 5’-CAGTCAAATCTGGTGGCATC-3’ human-*NQO1*-Forward 5’-AACTTTCAGAAGGGCCAGGT-3’ human-*NQO1*-Reverse 5’-CTGGGCTCTCCTTGTTGC-3’ human-*GPX4*-Forward 5’-TCGGCCGCCTTTGCC-3’ human-*GPX4*-Reverse 5’-TTCCCGAACTGGTTACACGG-3’ human-*GAPDH*-Forward 5’-CTGACTTCAACAGCGACACC-3’ human-*GAPDH*-Reverse 5’-TGCTGTAGCCAAATTCGTTGT-3’

### Organotypic brain slice cultures (OBSC)

OBSC’s were cultured as previously described [28]. Briefly, 6 month old *h*α*-Syn*^*-*^*/Nrf2*^*+/+*^, *h*_α_*-Syn*^*-*^*/Nrf2*^*-/-*^, *h*_α_*-Syn*^*+*^*/Nrf2*^*+/+*^, and *h*_α_*-Syn*^*+*^*/Nrf2*^*-/-*^ mice were rapidly decapitated and the brain placed in ice-cold Hanks’ balanced salt solution (HBSS, with Ca^2+^ and Mg^2+^; Life Technologies) supplemented with 25 mM HEPES (Life Technologies). The Compresstome® VF-300-0Z Vibrating Microtome was used to cut 350-μm-thick sections that were immediately plated on a hydrophilic PTFE cell culture insert (pore size: 0.4 μm; Millicell-CM, Millipore) in 50% DMEM (Life Technologies), 25% heat-inactivated horse serum (Invitrogen), 25% HBSS, 35 mM glucose (Sigma-Aldrich), and 25 mM HEPES. The sections were incubated at 37°C in a 5% CO_2_ atmosphere. After 1 day in vitro, the media was changed to 70% DMEM and 5% heat-inactivated horse serum. For treatment, slices were pre-incubated for 1 hr with DFO (100 µM) and then treated with erastin 10 µM for additional 16 hours. To assess ferroptotic death, OBSCs were incubated with 1 μg/ml propidium iodide (PI; Sigma-Aldrich) for 30 minutes. For immunostaining, treated and PI-stained OBSCs were, fixed in 4% paraformaldehyde for 2 hours, and then blocked in 10% goat serum for 1 hour at room temperature. Next, slices were incubated overnight at 4°C with anti-NeuN (1:500 – MAB377 Millipore, St. Louis, MO) primary antibody. After the sections were washed 3 times, they were incubated for 2 hours in Alexa Fluor 488–conjugated secondary antibody (Life Technologies-Molecular Probes, Grand Island, NY) at room temperature.

### Transmission Electron Microscopy

Brains were extracted and four regions, namely the midbrain, striatum, hippocampus, and cortex, were microdissected. Tissue slices for transmission (TEM) electron microscopy were fixed in 2.5% glutaraldehyde + 2% PFA in 0.1 M piperazine-N,N′-bis (2-ethanesulfonic acid) or PIPES buffer (pH 7.4) for 1 hr at RT or at 4°C overnight. Samples were post-fixed in 1% osmium tetroxide in PIPES buffer for 1 hr following a wash in 0.05 M PIPES buffer + 0.05 M Glycine and 2 × 10 min washes in 0.1 M PIPES. Following two more 10 min washes in deionized water (DIW), samples for TEM were block stained in 2% aqueous uranyl acetate, washed in DIW, dehydrated through a graded series of alcohols, infiltrated with 1:1 alcohol and Spurr’s resin overnight, and then embedded in 100% Spurr’s resin overnight at 60°C. Sections were viewed using a FEI Tecnai Spirit electron microscope (FEI Company, Hillsboro, OR) operated at 100 kV. 8-bit TIFF images were captured through an AMT 4 Mpixel camera. Morphometric measurements were conducted in digital images using Image J (NIH) software. Assessment of mitochondrial morphology was performed as described previously [6].

### Western Blotting

Brain tissue from the four specified regions was homogenized using the TissueLyser II (QIAGEN, Germantown, MD) in 10 vol. (w/v) of cold RIPA buffer (25 mM Tris-HCl, pH 7.6, 150 mM NaCl, 1% Triton X-100, 1% sodium deoxycholate, 0.1% SDS) supplemented with 1 mM phenylmethylsulfonyl fluoride (PMSF) and a phosphatase inhibitor cocktail (PIC). Tissues were then sonicated for 10 s to generate total cell lysates. The protein concentration in the different lysates was determined via the bicinchoninic acid method (BCA, Thermo/Pierce, Waltham, MA). Lysates were then boiled, sonicated, and resolved by SDS-PAGE, and membranes were subjected to appropriate antibodies at 4°C for overnight. Then membranes were incubated with anti-mouse or anti-rabbit horseradish peroxidase conjugated secondary antibodies (1:3000, Sigma Aldrich, St. Louis, MO) for 1 hr. All immunoblot images were taken using the Azure Biosystems 600 (Azure cSeries Advanced Imaging Systems, Dublin, CA). Relative densitometry analysis of western blots was performed using ImageJ Program (NIH). The primary antibodies applied were as follows: anti-NRF2 (1:1000, sc-13032), anti-FTH1 (1:1000, sc-376594), anti-GPx4 (1:1000, sc-166570), anti-AKR1C1 (1:1000, sc-398596), and anti-GCLM (1:1000, sc-166603) were from Santa Cruz Biotechnology (Dallas, TX). Anti-phospho Syn S129 (1:500, ab-59264), and anti-4-HNE (1:2000, ab-46545) were from Abcam (Cambridge, UK). Anti-α-Syn (1:1000, AHB0261) and anti-nitrotyrosine (1:500, A-21285) were from Life Technologies (Carlsbad, CA). α-Syn (clone 42, 1:1000, 610787) was from BD Biosciences (San Jose, CA). Anti-LC3 (1:2000, L7543) and anti-β-actin (1:5000, A2066) were from Sigma (St. Louis, MO). Anti-NCOA4 (ARA70, 1:2000) was from Bethyl Laboratories (Montgomery, TX). Anti-COX2 (1:1000, 27308-1-AP) and anti-ACSL4 (1:1000, 22401-1-AP) from Proteintech (Rosemont, IL). Anti-SLC7A11 (1:1000, CS #12691) was from Cell Signaling (Danvers, MA). Anti-SQSTM1 (p62, 1:2000, H00008878-M01) was from Abnova (Taipei, Taiwan).

### Statistical analysis

GraphPad Prism 8 software (San Diego, CA) was used for statistical analyses. For comparing two groups, unpaired t tests were used. For comparisons between three or more groups, one-way analysis of variance (ANOVA) followed by Tukey’s post-hoc test for multiple comparisons between treatment groups was conducted. For analyzing the nest building behavior, a two-way repeated measures ANOVA was performed using the Sigma Plot 14 (San Jose, CA) software. Differences were accepted as significant at p < 0.05. *p<0.05, **p<0.01.

## Acknowledgements

This study was funded by the following grants from the National Institutes of Health: P42ES004940 and R35ES031575 (DDZ)

## Notes

### Competing Interest Statement

The authors have declared no competing interest.

## References

[1] W. Yang, J.L. Hamilton, C. Kopil, J.C. Beck, C.M. Tanner, R.L. Albin, E. Ray Dorsey, N. Dahodwala, I. Cintina, P. Hogan, T. Thompson, Current and projected future economic burden of Parkinson’s disease in the U.S, NPJ Parkinsons Dis 6 (2020) 15.

[2] R.M. Meade, D.P. Fairlie, J.M. Mason, Alpha-synuclein structure and Parkinson’s disease - lessons and emerging principles, Mol Neurodegener 14(1) (2019) 29.

[3] C.R. Fields, N. Bengoa-Vergniory, R. Wade-Martins, Targeting Alpha-Synuclein as a Therapy for Parkinson’s Disease, Front Mol Neurosci 12 (2019) 299.

[4] M. Dodson, M.R. de la Vega, A.B. Cholanians, C.J. Schmidlin, E. Chapman, D.D. Zhang, Modulating NRF2 in Disease: Timing Is Everything, Annu Rev Pharmacol Toxicol 59 (2019) 555–575.

[5] M. Dodson, R. Castro-Portuguez, D.D. Zhang, NRF2 plays a critical role in mitigating lipid peroxidation and ferroptosis, Redox Biol (2019) 101107.

[6] S.J. Dixon, K.M. Lemberg, M.R. Lamprecht, R. Skouta, E.M. Zaitsev, C.E. Gleason, D.N. Patel, A.J. Bauer, A.M. Cantley, W.S. Yang, B. Morrison, 3rd, B.R. Stockwell, Ferroptosis: an iron-dependent form of nonapoptotic cell death, Cell 149(5) (2012) 1060–72.

[7] J.H. Suh, S.V. Shenvi, B.M. Dixon, H. Liu, A.K. Jaiswal, R.M. Liu, T.M. Hagen, Decline in transcriptional activity of Nrf2 causes age-related loss of glutathione synthesis, which is reversible with lipoic acid, Proc Natl Acad Sci U S A 101(10) (2004) 3381–6.

[8] A. Cuadrado, Brain-Protective Mechanisms of Transcription Factor NRF2: Toward a Common Strategy for Neurodegenerative Diseases, Annu Rev Pharmacol Toxicol 62 (2022) 255–277.

[9] A. Anandhan, N. Nguyen, A. Syal, L.A. Dreher, M. Dodson, D.D. Zhang, L. Madhavan, NRF2 Loss Accentuates Parkinsonian Pathology and Behavioral Dysfunction in Human alpha-Synuclein Overexpressing Mice, Aging Dis 12(4) (2021) 964–982.

[10] E. Deas, N. Cremades, P.R. Angelova, M.H. Ludtmann, Z. Yao, S. Chen, M.H. Horrocks, B. Banushi, D. Little, M.J. Devine, P. Gissen, D. Klenerman, C.M. Dobson, N.W. Wood, S. Gandhi, A.Y. Abramov, Alpha-Synuclein Oligomers Interact with Metal Ions to Induce Oxidative Stress and Neuronal Death in Parkinson’s Disease, Antioxid Redox Signal 24(7) (2016) 376–91.

[11] P.R. Angelova, M.L. Choi, A.V. Berezhnov, M.H. Horrocks, C.D. Hughes, S. De, M. Rodrigues, R. Yapom, D. Little, K.S. Dolt, T. Kunath, M.J. Devine, P. Gissen, M.S. Shchepinov, S. Sylantyev, E.V. Pavlov, D. Klenerman, A.Y. Abramov, S. Gandhi, Alpha synuclein aggregation drives ferroptosis: an interplay of iron, calcium and lipid peroxidation, Cell Death Differ 27(10) (2020) 2781–2796.

[12] S. Masaldan, A.I. Bush, D. Devos, A.S. Rolland, C. Moreau, Striking while the iron is hot: Iron metabolism and ferroptosis in neurodegeneration, Free Radic Biol Med 133 (2019) 221–233.

[13] L. Ma, M. Gholam Azad, M. Dharmasivam, V. Richardson, R.J. Quinn, Y. Feng, D.L. Pountney, K.F. Tonissen, G.D. Mellick, I. Yanatori, D.R. Richardson, Parkinson’s disease: Alterations in iron and redox biology as a key to unlock therapeutic strategies, Redox Biol 41 (2021) 101896.

[14] B.G. Trist, D.J. Hare, K.L. Double, Oxidative stress in the aging substantia nigra and the etiology of Parkinson’s disease, Aging Cell 18(6) (2019) e13031.

[15] P.R. Angelova, N. Esteras, A.Y. Abramov, Mitochondria and lipid peroxidation in the mechanism of neurodegeneration: Finding ways for prevention, Med Res Rev 41(2) (2021) 770–784.

[16] A. Anandhan, M. Dodson, C.J. Schmidlin, P. Liu, D.D. Zhang, Breakdown of an Ironclad Defense System: The Critical Role of NRF2 in Mediating Ferroptosis, Cell Chem Biol 27(4) (2020) 436–447.

[17] D. Beraud, H.A. Hathaway, J. Trecki, S. Chasovskikh, D.A. Johnson, J.A. Johnson, H.J. Federoff, M. Shimoji, T.R. Mhyre, K.A. Maguire-Zeiss, Microglial activation and antioxidant responses induced by the Parkinson’s disease protein alpha-synuclein, J Neuroimmune Pharmacol 8(1) (2013) 94–117.

[18] L. Gan, M.R. Vargas, D.A. Johnson, J.A. Johnson, Astrocyte-specific overexpression of Nrf2 delays motor pathology and synuclein aggregation throughout the CNS in the alpha-synuclein mutant (A53T) mouse model, J Neurosci 32(49) (2012) 17775–87.

[19] J.A. Johnson, D.A. Johnson, A.D. Kraft, M.J. Calkins, R.J. Jakel, M.R. Vargas, P.C. Chen, The Nrf2-ARE pathway: an indicator and modulator of oxidative stress in neurodegeneration, Ann N Y Acad Sci 1147 (2008) 61–9.

[20] I. Lastres-Becker, A. Ulusoy, N.G. Innamorato, G. Sahin, A. Rabano, D. Kirik, A. Cuadrado, alpha-Synuclein expression and Nrf2 deficiency cooperate to aggravate protein aggregation, neuronal death and inflammation in early-stage Parkinson’s disease, Hum Mol Genet 21(14) (2012) 3173–92.

[21] V. Vasquez, J. Mitra, G. Perry, K.S. Rao, M.L. Hegde, An Inducible Alpha-Synuclein Expressing Neuronal Cell Line Model for Parkinson’s Disease1, J Alzheimers Dis 66(2) (2018) 453–460.

[22] F. Liu, J. Hindupur, J.L. Nguyen, K.J. Ruf, J. Zhu, J.L. Schieler, C.C. Bonham, K.V. Wood, V.J. Davisson, J.C. Rochet, Methionine sulfoxide reductase A protects dopaminergic cells from Parkinson’s disease-related insults, Free Radic Biol Med 45(3) (2008) 242–55.

[23] I. Barde, P. Salmon, D. Trono, Production and titration of lentiviral vectors, Curr Protoc Neurosci Chapter 4 (2010) Unit 4 21.

[24] M.F. Chesselet, F. Richter, C. Zhu, I. Magen, M.B. Watson, S.R. Subramaniam, A progressive mouse model of Parkinson’s disease: the Thy1-aSyn (“Line 61”) mice, Neurotherapeutics 9(2) (2012) 297–314.

[25] S.M. Fleming, J. Salcedo, P.O. Fernagut, E. Rockenstein, E. Masliah, M.S. Levine, M.F. Chesselet, Early and progressive sensorimotor anomalies in mice overexpressing wild-type human alpha-synuclein, J Neurosci 24(42) (2004) 9434–40.

[26] H.A. Lam, N. Wu, I. Cely, R.L. Kelly, S. Hean, F. Richter, I. Magen, C. Cepeda, L.C. Ackerson, W. Walwyn, E. Masliah, M.F. Chesselet, M.S. Levine, N.T. Maidment, Elevated tonic extracellular dopamine concentration and altered dopamine modulation of synaptic activity precede dopamine loss in the striatum of mice overexpressing human alpha-synuclein, J Neurosci Res 89(7) (2011) 1091–102.

[27] M.J. Corenblum, A.J. Flores, M. Badowski, D.T. Harris, L. Madhavan, Systemic human CD34(+) cells populate the brain and activate host mechanisms to counteract nigrostriatal degeneration, Regen Med 10(5) (2015) 563–77.

[28] Q. Li, X. Han, X. Lan, Y. Gao, J. Wan, F. Durham, T. Cheng, J. Yang, Z. Wang, C. Jiang, M. Ying, R.C. Koehler, B.R. Stockwell, J. Wang, Inhibition of neuronal ferroptosis protects hemorrhagic brain, JCI Insight 2(7) (2017) e90777.

