## Supplemental Figures 1 and 2 for "α-syn overexpression, NRF2 suppression, and enhanced ferroptosis create a vicious cycle of neuronal loss in Parkinson’s disease"

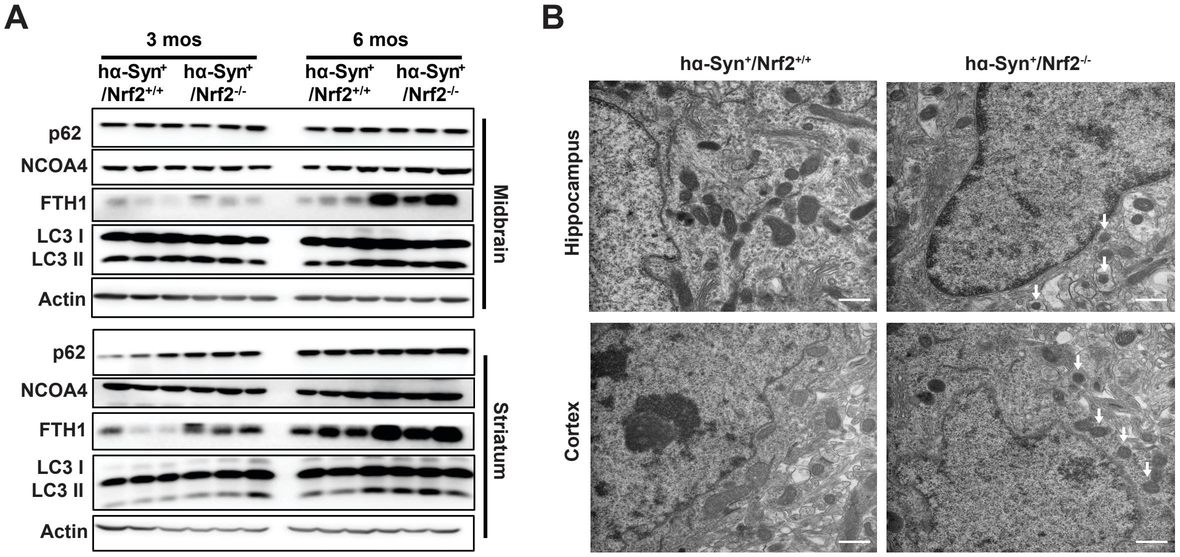


**Figure S1: Mitochondrial morphology and ferritinophagy are altered in 6-month-old *hα-Syn*$^{+}$*:Nrf2*$^{-/-}$ mice.** (A) TEM micrographs of the hippocampus and cortex of 6-month-old *hα-Syn*$^{+}$*:Nrf2*$^{+/+}$ and *hα-Syn*$^{+}$*:Nrf2*$^{-/-}$ mice. Arrows indicate condensed mitochondria. Scale bar = 500 nm. (B) Immunoblot analysis of p62, Fth1, and LC3-I/II protein levels from the MB and ST of 3 and 6 month old *hα-Syn*$^{+}$*:Nrf2*$^{+/+}$ and *hα-Syn*$^{+}$*:Nrf2*$^{-/-}$ mice. β-actin was used as an internal loading control.

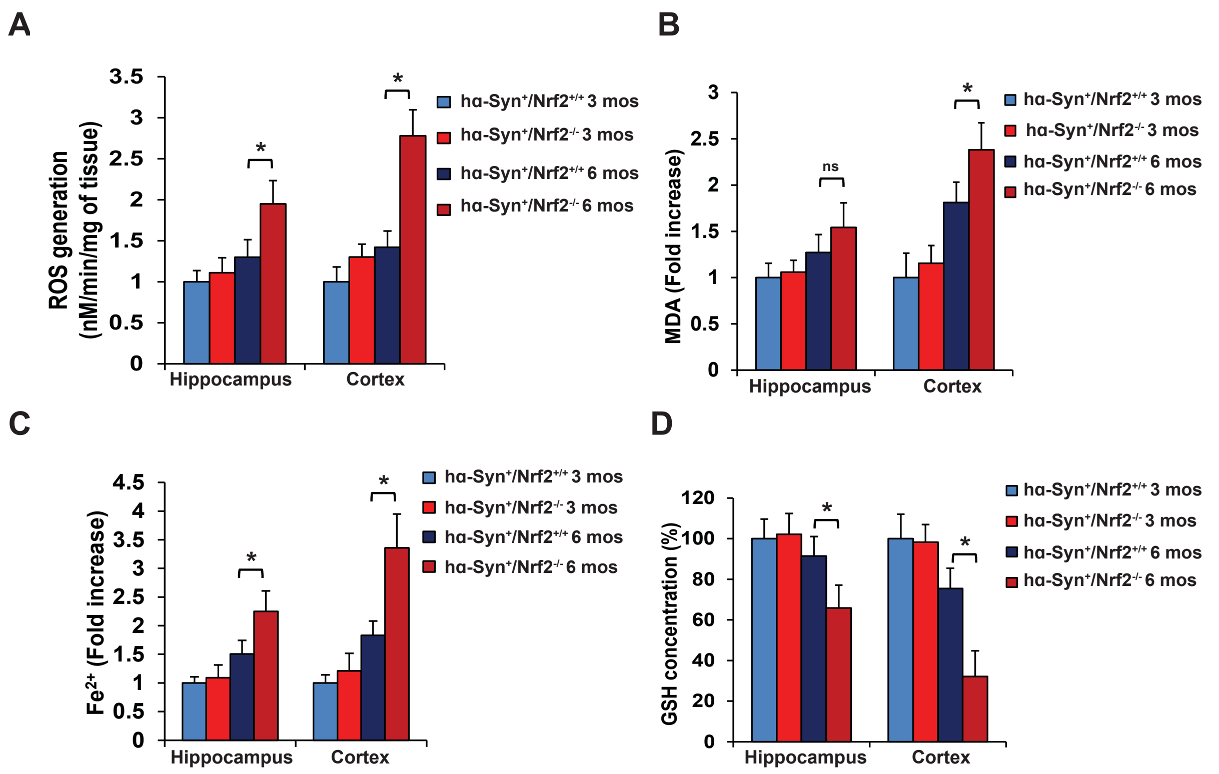


**Figure S2: ROS, lipid peroxidation, free iron, and GSH levels in the hippocampus and cortex of 3 and 6 month old *hα-Syn$^{+}$:Nrf2$^{+/+}$* and *hα-Syn$^{+}$:Nrf2$^{-/-}$* mice.** The hippocampus and cortex of 3 and 6 month old *hα-Syn$^{+}$:Nrf2$^{+/+}$* and *hα-Syn$^{+}$:Nrf2$^{-/-}$* mice were collected and assessed for changes related to loss of NRF2. (A) ROS generation measured by electron paramagnetic resonance (EPR) spectroscopy. n=6 (B) Malondialdehyde (MDA) formation measured by TBARs assay. n=6 (C) Free ferrous iron (Fe2+) measured via Ferene-S based colorimetry. n=6 (D) Total glutathione (GSH) levels measured via Quantichrom assay. *$p<0.05$, One-way ANOVA with Tukey's post-hoc test.
